# Living in a changing world: effects of roads and *Pinus* monocultures on an anuran metacommunity in southern Brazil

**DOI:** 10.1101/2020.07.31.231647

**Authors:** Diego Anderson Dalmolin, Alexandro Marques Tozetti, Maria João Ramos Pereira

## Abstract

Amphibians are undergoing global-scale declines due to the increased incidence of anthropogenic stressors. The loss of species with unique evolutionary histories and functional traits poses a serious risk to the maintenance of ecosystem functions in aquatic environments, already directly affected by several anthropogenic land-use changes. Here, we investigated the influence of anthropogenic stressors (roads and *Pinus* monocultures) on functional, phylogenetic and taxonomic composition and functional dispersion of an anuran metacommunity of 33 ponds in southern Brazil. We expected for the relative influence of anthropogenic stressors to vary according to the compositional facet, with a greater influence of these stressors on the functional and phylogenetic than on the taxonomic facet. We also expected traits related to habitat exploration (head shape and eye size and position) to be more influenced by *Pinus* monocultures, while the traits related to the dispersion and the physiological control of individuals (limb length and body mass) to be similarly influenced by roads and *Pinus*. To evaluate this, we used PERMANOVA analyses for each of the compositional facets and anthropogenic stressor, and path models to verify all possible relationships between patterns of functional dispersion and anthropogenic stressors. We found that, while the distance from ponds to *Pinus* monocultures influences the phylogenetic composition, distance to roads influences the functional composition; distance to roads affects mostly the functional dispersion of the communities. These anthropogenic stressors affect the structure of anuran communities, even those formed by generalist species in terms of habitat use. There is a decline in diversity in communities located close to *Pinus* and roads, leading to losses in the evolutionary history accumulated in these communities. The control of vehicle traffic during reproduction periods and the maintenance of areas with natural vegetation, particularly around ponds, may help mitigate the negative effects of anthropogenic stressors on anuran communities.

## Introduction

The transformation of natural habitats into anthropogenic landscapes is considered one of the major threats to biodiversity in the 21^st^ century (Bitar et al., 2014). Habitat loss and fragmentation resulting from agriculture, monoculture forestry and urban expansion results in changes in the structure and functioning of ecosystems throughout the planet (Lion, Garda & Fonseca, 2014; Costa, Solé & Nomura, 2017; Berriozabal-Islas et al., 2018). At this present age, known as the Anthropocene, attention to the effects of anthropogenic changes on all levels and dimensions of biological diversity has become indispensable (Shanafelt et al., 2018).

The conservation of species with complex life-histories, such as anurans, is challenging (Salice et al., 2011). Anuran life-cycle alternates between an aquatic or semi-aquatic larval phase and an adult able to move in land and to transition between aquatic and terrestrial environments. These characteristics of the anuran life-cycle make them particularly vulnerable to anthropogenic modifications. Increased levels of ultraviolet (UV) radiation (Hite et al., 2016), pollution and the introduction of exotic species (Both et al., 2014; Both & Melo, 2015) are associated with the increased incidence of deleterious and lethal pathogens in anurans. The transformation of natural areas into monocultures and the expansion of the road network drastically modify the structure of the landscapes; consequently, species behaviour, population dynamics and gene flow between populations may be severely affected (Bitar et al., 2014; Berriozabal-Islas et al., 2018).

Together, the aforementioned changes tend to alter species’ behavioural patterns (especially those associated with the reproductive repertoire, such as advertisement calls and the choice of breeding sites), to increase mortality rates (Lips et al., 2006), and to decrease recruitment (Hayes et al., 2010). Consequently, population declines, as well as reductions in genetic diversity within and between populations, have been reported for several anuran species in human-modified areas (Pineda et al., 2005; Lion et al., 2014; Berriozabal-Islas et al., 2018). However, some studies reported increases in anuran diversity in environments with moderate levels of anthropogenic changes (Wanger et al., 2010; Bitar et al., 2014; Pelinson, Garey & Rossa-Feres, 2016), eventually supporting the intermediate disturbance hypothesis (Wilkinson, 1999) and demonstrating that, at least to a certain degree, anthropogenic disturbance is not always associated with biodiversity loss (Berriozabal-Islas et al., 2018).

Many ecological models neglect the effect of anthropogenic changes in the landscape on biological communities (Schmitz, 2016; Shanafelt et al., 2018). Additionally, ecological interpretations are mostly based on taxonomic information, ignoring eco-evo relationships (Webb et al., 2002; Sobral & Cianciaruso, 2016; Arnan, Cerdá & Retana, 2017). Ignoring potential distinct responses of taxonomic, functional and phylogenetic compositions is a major shortcoming in ecological studies, as human activities may cause severe changes in composition by eliminating unique functional traits (Tilman, 2001) or unique evolutionary lineages (Magurran, 2004). Such changes in functional and phylogenetic composition of communities may drastically alter ecosystem balance and the relative importance of different ecological processes (Hof, Rahbek & Araújo, 2010) resulting in short-term changes in biodiversity (Alberti, 2015). Anthropogenic stressors have already been identified as some of the main factors responsible for significant changes in the patterns of intraspecific variation of functional traits related to the dispersion and vertical exploration of habitat in anurans (Dalmolin, Tozetti & Ramos Pereira, 2020). Thus, the inclusion of functional and phylogenetic data, as well as taxonomic information, potentially leads to more accurate assessments of the actual conservation status of ecosystems facing anthropogenic threats (Webb et al., 2002). Previous studies have shown that, although anthropogenic disturbances may not immediately cause a steep decline in species richness, they may cause drastic reductions in functional and phylogenetic compositions and, in particular, in the average values of some traits of different groups of organisms, compromising the ecological functions they perform and threatening the resilience of ecosystems (Biswas & Mallik, 2010; Carreño & Rocabado et al., 2012; Ribeiro et al., 2017; Matuoka et al., 2020).

In this study we investigate the effects of potential anthropogenic stressors – roads and *Pinus* monocultures – on the composition of anuran communities in the Lagoa do Peixe National Park (PNLP), one of the two Ramsar sites in southern Brazil. Roads and *Pinus* monocultures were selected because they reflect the main anthropogenic changes in the region’s landscape and have known effects on community structure and acoustic behaviour of anurans (particularly traffic), potentially affecting the reproductive patterns of the species of this taxon (e.g. Saccol, Bolzan & Santos, 2017; Caorsi et al., 2017).

The distribution of anurans is conditioned by the adequacy of their morphological characteristics and physiological functions to environmental conditions and, mostly, by their dispersion abilities, which may be limited for some groups (Semlitsch, 2008; Oliveira et al., 2016). Many of these traits are phylogenetically conserved in anurans (Lourenço-de-Moraes et al., 2019). However, functional, phylogenetic and taxonomic structures may follow distinct patterns resulting from variation in composition (Ouchi-Melo et al., 2018; Dalmolin, Tozetti & Ramos Pereira, 2019). We thus expect for the relative influence of each anthropogenic stressor to vary with each compositional facet – taxonomic, functional, phylogenetic (Ouchi-Melo et al., 2018). We expect greater effects of anthropogenic variables on phylogenetic and functional composition. Indeed, up to a certain level, environmental disturbance allows for the coexistence between dominant competitors and fast colonizers (Chesson & Huntly, 1997; Roxburgh, Shea & Wilson, 2004), favouring the coexistence of a larger number of evolutionary lineages (Yuan et al., 2016). We expect patterns of functional dispersion to be associated to different stressors: functional traits related to habitat exploration (head shape and eye size and position) to be more influenced by stressors related to the monocultures, while traits related to dispersion and physiological control of individuals (relative limb length and body mass) to be similarly influenced by both roads and *Pinus* monocultures.

## Material and Methods

### Ethics statement

We obtained sampling permits from Instituto Chico Mendes de Conservação da Biodiversidade (ICMBio) (licence 55409). Our sampling did not involve any endangered or protected species. We restricted amphibian manipulation in the field to the minimum necessary; specimens collected were identified to the species level, measured and immediately released after these procedures in the same pond/site where they were captured.

### Study Area

The Lagoa do Peixe National Park (PNLP; 31°02 – 31°48 S; 50°77 – 51°15 W; figure 1a-e) comprises over 34,000 hectares of protected wetlands. PNLP has 64 km length and 6 km width, and integrates one of the regions of southern Brazil with higher concentration of wetlands: the coastal plain of the state of Rio Grande do Sul RAMSAR (2018). PNLP presents subtropical humid climate, and temperatures range between 13 °C and 24 °C with annual average of 17.5 °C. The mean annual precipitation varies between 1200 and 1500 mm (Maltchik et al., 2003).

**Fig. 1.**
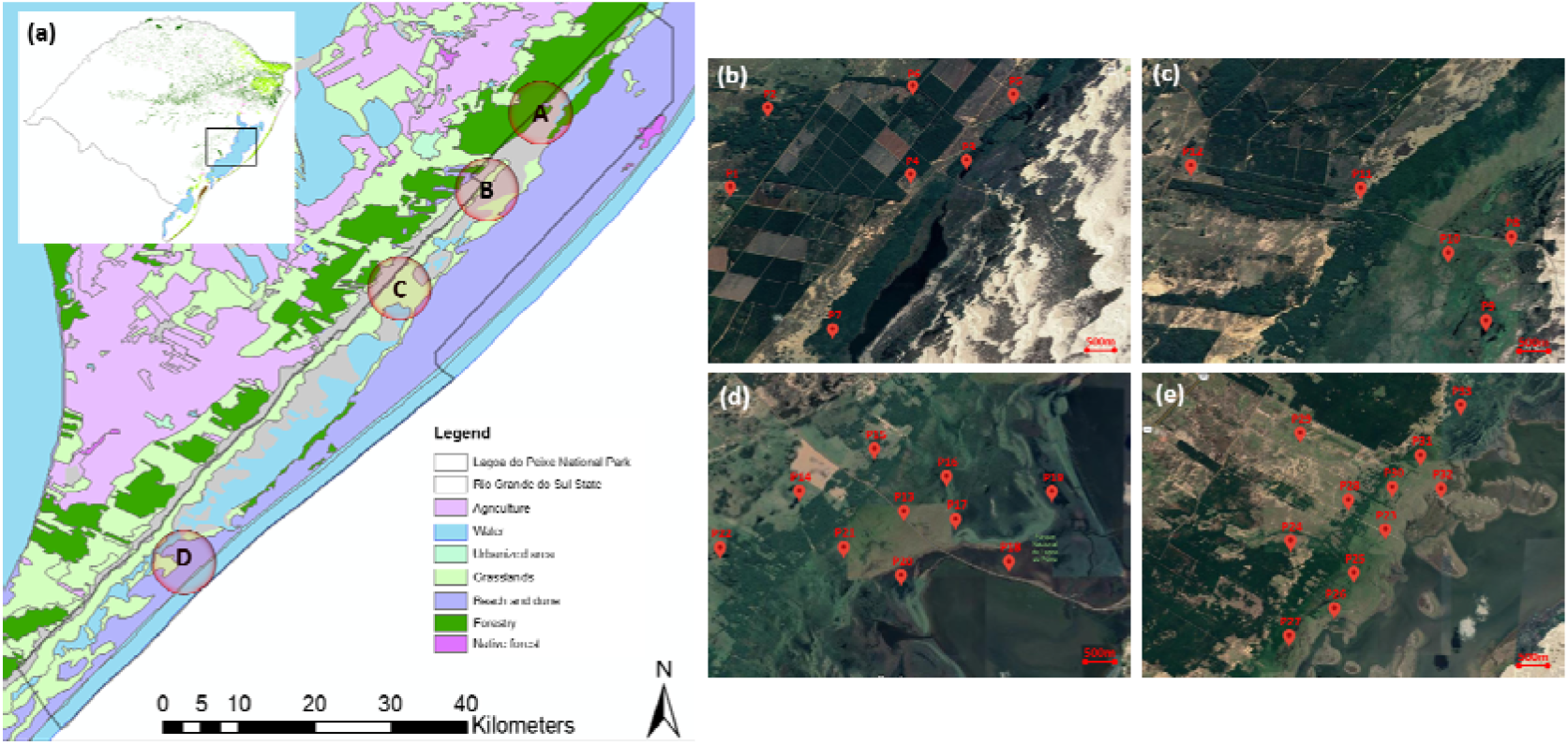
Location of the study area in Rio Grande do Sul, southern Brazil. Land use (a) and study areas (b-e) at Lagoa do Peixe National Park. The sampled ponds are represented by the red icons (N = 33).

### Anuran surveys, trait measurement and phylogenetic hypothesis

From October 2016 to March 2017 we sampled adult anurans in 33 ponds throughout the study region (Fig. 1). We focused on adults because they seem to be more susceptible to the effects of anthropogenic stressors due to their dispersal between ponds. We selected ponds according their biotic and abiotic characteristics (see Table S2), spatial independence, accessibility and landowner permission. Distance between ponds ranged from 0.7 to 39 km. We used calling surveys and active searches at breeding habitats to find adult anurans in each pond (Scott & Woodward, 1994). For the two search types, the perimeter of the ponds was covered by two researchers (D.A.D. and a field assistant); all adult anurans found and captured were immediately identified and measured. The surveys were carried out from 6 p.m. to 0 a.m, totalling 6 hours of sampling per pond.

We measured five morphological traits in each individual captured: head shape; eye position; eye size; relative limb length; and body mass (Table 1; Figure 2). Additionally, we compiled six additional life-history traits from the available literature: reproductive mode; number of eggs per reproductive event; activity period; preferred habitat type; presence/absence of fossorial habits; and reproductive season. All attributes were used to build a pairwise distance matrix between species, using the Gower standardization for mixed variables (Legendre & Legendre, 2012).

**Table 1:**
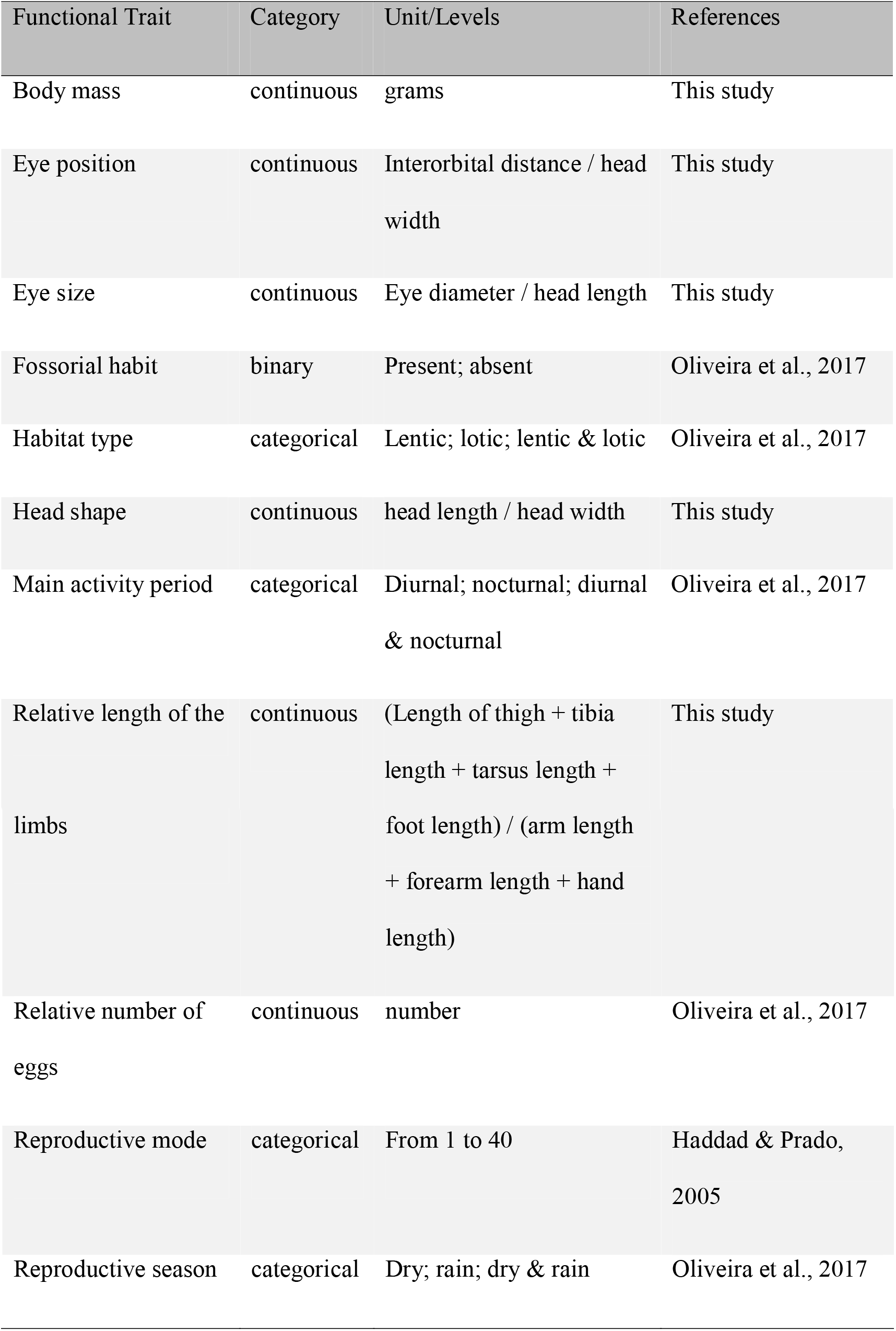
Anuran functional traits.

**Fig. 2.**
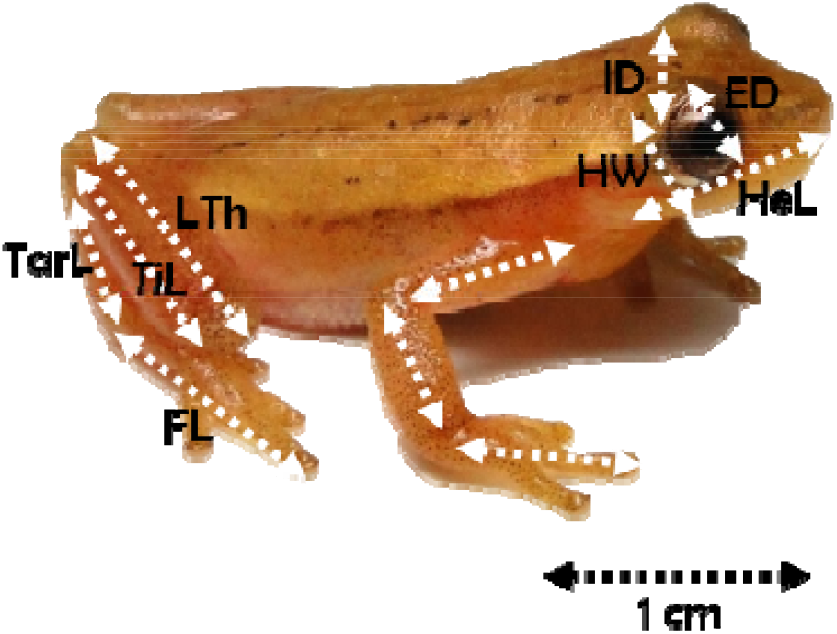
Morphological metrics evaluated in adults (except body mass): length of thigh (LTh); tibia length (TiL); tarsus length (TarL); foot length (FL); arm length (AL); forearm length (FL); hand length (HL); head length (HeL); head width (HW); interorbital distance (ID); eye diameter (ED). In this picture: a male of *Dendropsophus sanborni* (Amphibia, Hylidae) captured at the Parque Nacional da Lagoa do Peixe, Brazil.

Finally, we built a phylogenetic tree by pruning the amphibian phylogeny proposed by Jetz and Pyron (2018) to include only the species found in the whole of the sampled ponds using the function *prune.sample* of the R package picante (Kembel et al., 2010). Then a matrix of phylogenetic distances between the species occurring in the ponds was built.

### Anthropogenic stressors, local environmental and spatial variables

The anthropogenic descriptors evaluated and their description are presented in Table 2. These variables included the total area occupied by roads and *Pinus* monocultures within a buffer of 1 km^2^ around the sampled pond, as well as the distance of each pond to the nearest road and *Pinus* plantation. These values were obtained using high-resolution aerial photographs available from Google Earth (http://earth.google.com/).

**Table 2:** Anthropogenic stressors measured between October 2016 and March 2017 at Lagoa do Peixe Nation Park, Rio Grande do Sul, Brazil.

| Descriptors | Description/levels | Ecological Relevance |
| --- | --- | --- |
| Distance to the nearest *Pinus* monoculture | Distance to the nearest pine monoculture (m) | May affect the physiological control and survival of individuals |
| Area occupied by *Pinus* monocultures | Area (m<sup>2</sup>) |  |
| Distance to the nearest road | Distance to the nearest pine monoculture (m) | May affect the dispersion, reproduction rates and survival of individuals |
| Area occupied by roads | Area (m<sup>2</sup>) |  |

### Data Analysis

Figure 3 summarizes the sequence of the analytical procedures employed.

**Fig. 3.**
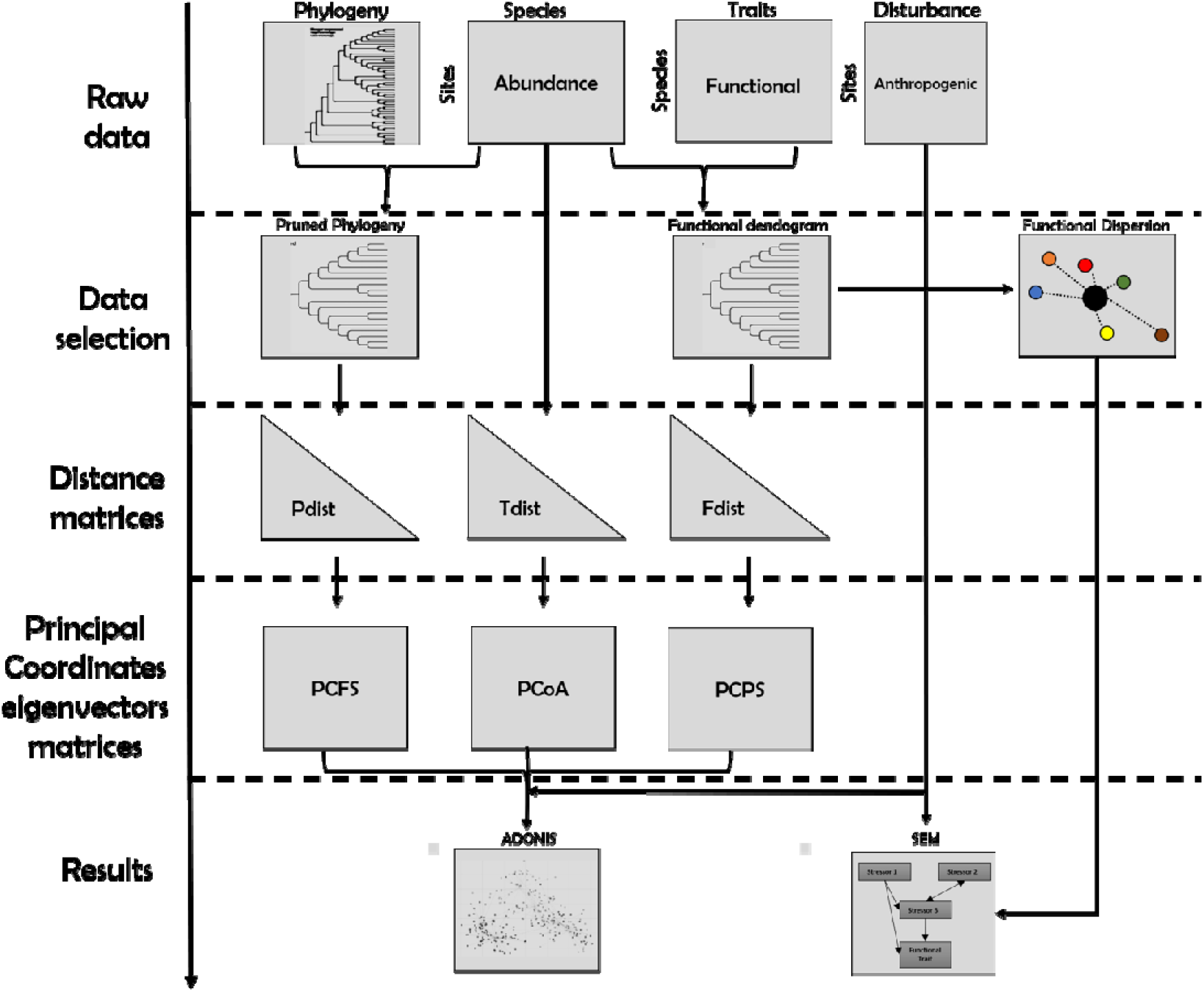
Flow chart of the analytical approach followed. For detailed descriptions and procedures, see the main text. Abbreviations: Fdist, Pdist and Tdist represent functional, phylogenetic and taxonomic compositional dissimilarity matrices; PCFS, PCPS and PCoA represent “eigenvectors” matrices of the principal components of functional, phylogenetic and taxonomic structure analysis, respectively.

### Data matrices

Initially we created an abundance matrix containing the total species count for each pond in the month of highest recorded abundance. This procedure prevents underestimating population abundance caused by calculating the mean of successive samples and prevents overestimating caused by the re-counting of individuals if successive samples are summed up (Scott & Woodward, 1994). We used the Hellinger distance to transform the abundance data; this procedure homogenizes variation between species’ abundances (Legendre & Legendre, 2012). We then created a trait matrix containing the average trait values for each species in the abundance matrix. Following, we created an anthropogenic stressor matrix, containing area and distance to roads and pine monocultures relative to each pond. We standardized the values of the anthropogenic stressors by subtracting each value from the average of the corresponding variable and dividing the result by its standard deviation.

### Anuran functional, phylogenetic and taxonomic composition

We used a principal coordinate of phylogenetic structure analysis (PCPS) to assess the relationship of all clades occurring in the ponds (Duarte, 2011). This analysis results in vector ordination expressing orthogonal gradients in phylogenetic composition across communities (Duarte, 2011; Carlucci et al., 2017), thus allowing the identification of lineages that better represent different parts of anthropogenic, local environmental and spatial gradients (Duarte et al., 2014; Carlucci et al., 2017). PCPS analysis is built according to the phylogenetic distance between each pair of species recorded in the communities. Finally, the correlation between PCPS vectors and species is used to evaluate the lineage commonness across communities (Duarte, 2011; Carlucci et al., 2017). This analysis was done using the function “pcps” of the PCPS R package. We used this same approach for the functional composition. First, we built a functional dendrogram with the information present in the abundance and the functional matrices. Then, we used the functional distance between each pair of species recorded in the communities to build the vectors of the principal coordinates of functional structure (in this case, PCFS; Pillar & Duarte, 2010). Finally, we assessed the taxonomic composition by using principal coordinates analysis (PCoA) in the ape R package. Similar to the PCPS, the “pcoa” function takes into account the matrix of taxonomic distance between each pair of species to compute the principal coordinate decomposition (Gower, 1966).

### Effects of anthropogenic stressors on anuran composition

We assessed the relative effects of each anthropogenic stressor on anuran functional, phylogenetic and taxonomic composition by using permutational multivariate analysis of variance with 9999 random permutations (PerMANOVA, ‘adonis’ function in vegan R package; Oksanen et al., 2014); we ran this analysis separately for each of the compositional dimensions. The variance explained by each fraction was based on the adjusted *R^2^* (Blanchet, Legendre & Bocard, 2008).

### Path models

We developed a theoretical model of how the different anthropogenic stressors may affect the patterns of functional dispersion of the five functional traits evaluated (Fig. 4). We used structural equation modelling (SEM; Fox, 2010) to examine the support for each model for each functional trait analysed and for the set of all functional traits. We used the lavaan package (Rosseel, 2012) in R (RStudio Team, 2020) for SEM. We assessed the goodness-of-fit of each model using the chi-square test (χ^2^): models with low χ^2^ and non-significant P-values (P > 0.05) were considered as good models because they indicate consistency between observed data and the hypothesised model (Grace, 2006). We expressed the results using graphical models, where the arrows pointing from the putative “cause” to the “effect” represent the causation hypotheses.

**Fig 4.**
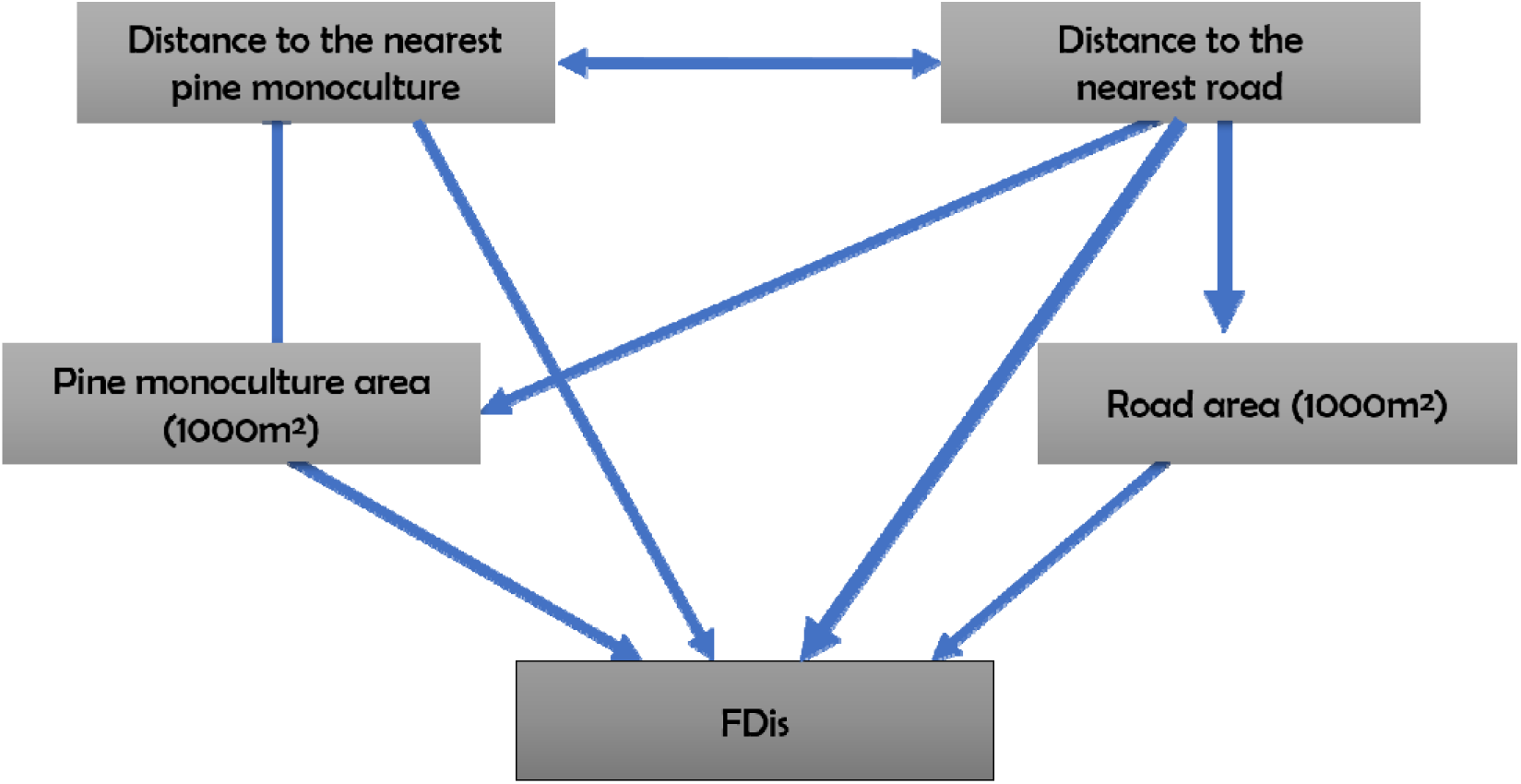
Theoretical causal model explaining the relationships between functional traits associated to dispersal in anurans and the potential underlying anthropogenic stressors. FDis = Functional dissimilarity matrices.

## Results

### Anuran richness and abundance

Eleven species of anurans belonging to three families (Bufonidae, Hylidae and Leptodactylidae) were registered. The most frequent species were *Dendropsophus sanborni* and *Pseudis minuta,* occurring in 19 of the 33 ponds evaluated. *Physalaemus biligonigerus* and *Scinax fuscovarius* were less frequent occurring, respectively, in three and four of the sampled ponds (for the complete list of species and occurrence patterns in the sampled ponds see Table S1).

### Predictors of compositional dimensions

Functional composition was mostly affected by the distance to roads, while phylogenetic composition was mostly influenced by the distance to *Pinus* monocultures (Table 3). Taxonomic composition was not significantly affected by any of the anthropogenic stressors evaluated here.

**Table 3:** Effects of anthropogenic stressors on functional, phylogenetic and taxonomic composition of an anuran metacommunity accounted by PerMANOVA. Bold values represent statistically significant effect values.

| Anthropogenic stressors | Sum of squares | **R<sup>2</sup>** | **F** | ***p*** |
| --- | --- | --- | --- | --- |

| Functional composition |  |  |  |  |
| --- | --- | --- | --- | --- |
| Distance to the nearest pine monoculture | 0.03 | 0.04 | 0.94 | 0.41 |
| Pine monoculture area (1000 m<sup>2</sup>) | 0.04 | 0.05 | 1.26 | 0.24 |
| Distance to the nearest road | 0.10 | 0.11 | 2.86 | **0.03** |
| Road area (1000 m<sup>2</sup>) | 0.03 | 0.04 | 0.96 | 0.39 |
| Phylogenetic composition |  |  |  |  |
| Distance to the nearest pine monoculture | 0.30 | 0.09 | 2.41 | **0.04** |
| Pine monoculture area (1000 m<sup>2</sup>) | 0.26 | 0.06 | 1.61 | 0.14 |
| Distance to the nearest road | 0.28 | 0.07 | 1.69 | 0.11 |
| Road area (1000 m<sup>2</sup>) | 0.11 | 0.03 | 0.66 | 0.68 |
| Taxonomic composition |  |  |  |  |
| Distance to the nearest pine monoculture | 2.82 | 0.06 | 1.57 | 0.13 |
| Pine monoculture area (1000 m<sup>2</sup>) | 1.92 | 0.04 | 1.07 | 0.35 |
| Distance to the nearest road | 2.66 | 0.06 | 1.48 | 0.16 |
| Road area (1000 m<sup>2</sup>) | 1.99 | 0.04 | 1.11 | 0.36 |

### The influence of anthropogenic stressors on functional dispersion

In general, the average and the maximum values of functional dispersion in the set of communities evaluated here were similar between each of the functional traits and the set of all functional traits (Fig. 5). However, the body mass and the relative length of the limbs were the functional traits with lowest mean values of functional dispersion.

**Fig. 5.**
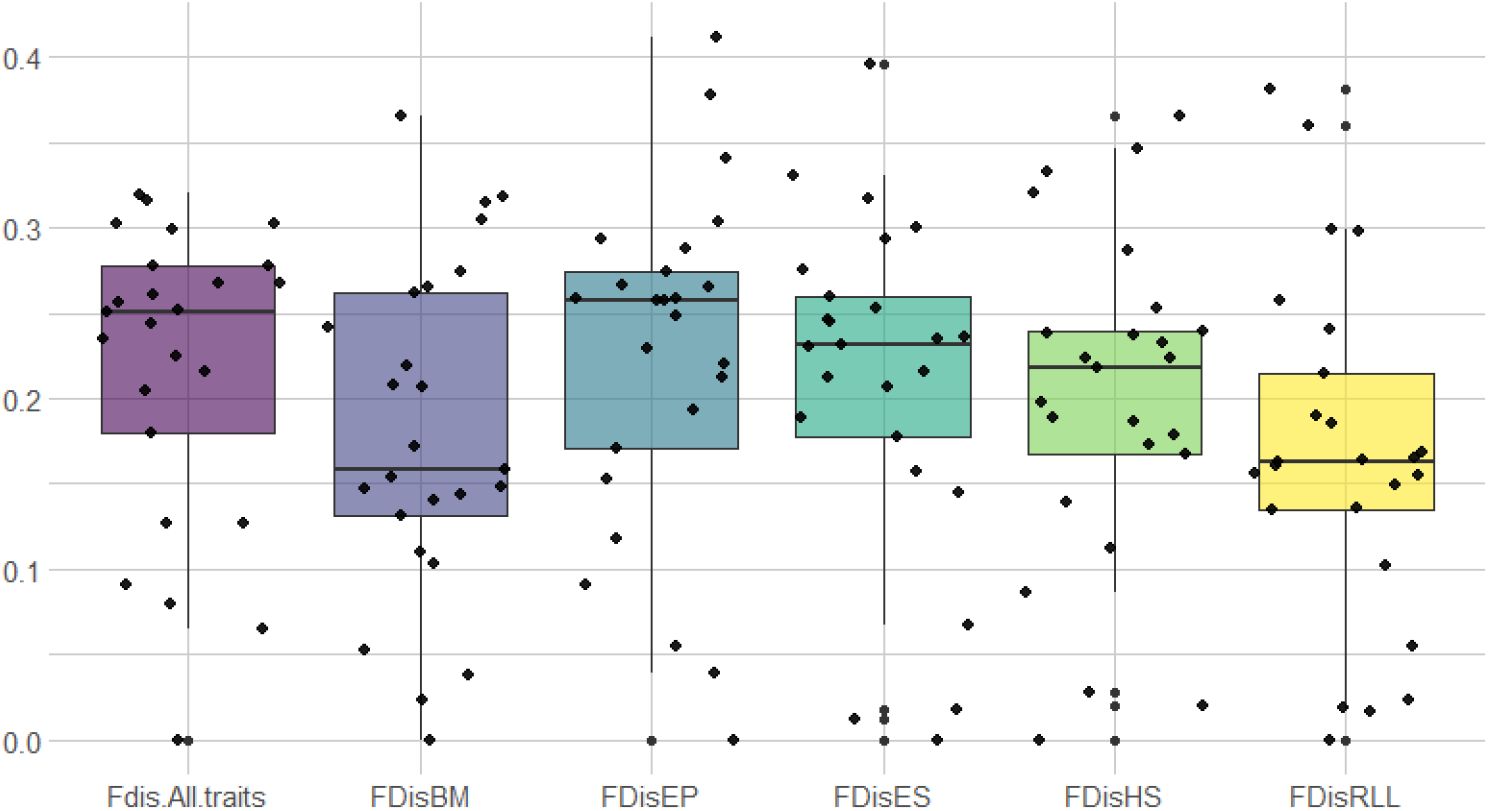
Box-plots showing the values of functional dispersion of each of the functional traits evaluated and the set comprising all the evaluated traits of the anuran metacommunity in Lagoa do Peixe National Park, southern Brazil. Black dots represent functional dispersion values for each of the communities. **Fdis.All.traits** - functional dispersion for all functional attributes evaluated; **FDisBM** - functional dispersion of body mass; **FDisEP** - functional dispersion of eyes position; **FDisES** - functional dispersion of eye size; **FDisHS** - functional dispersion of head shape; **FDisRLL** - functional dispersion of the relative length of the limbs.

We found support for the six tested path models (χ^2^ = 0.33; **P** = 0.85), suggesting that the evaluated anthropogenic stressors shape patterns of functional dispersion via different pathways of environmental modifications in the study area. In general, our models were able to explain over 60% of the observed variation in the patterns of functional dispersion; indeed, the model built for the relative length of the limbs presented the largest fraction explained for the variation observed (**R^2^** = 0.79). We also found that the patterns of functional dispersion of all of the evaluated traits was directly affected by the distance to the nearest road, similar to what was found for the functional composition (Fig. 6 a-f). However, this effect was not significant for body mass. We found no significant effects of the remaining anthropogenic variables on the patterns of functional dispersion.

**Fig. 6.**
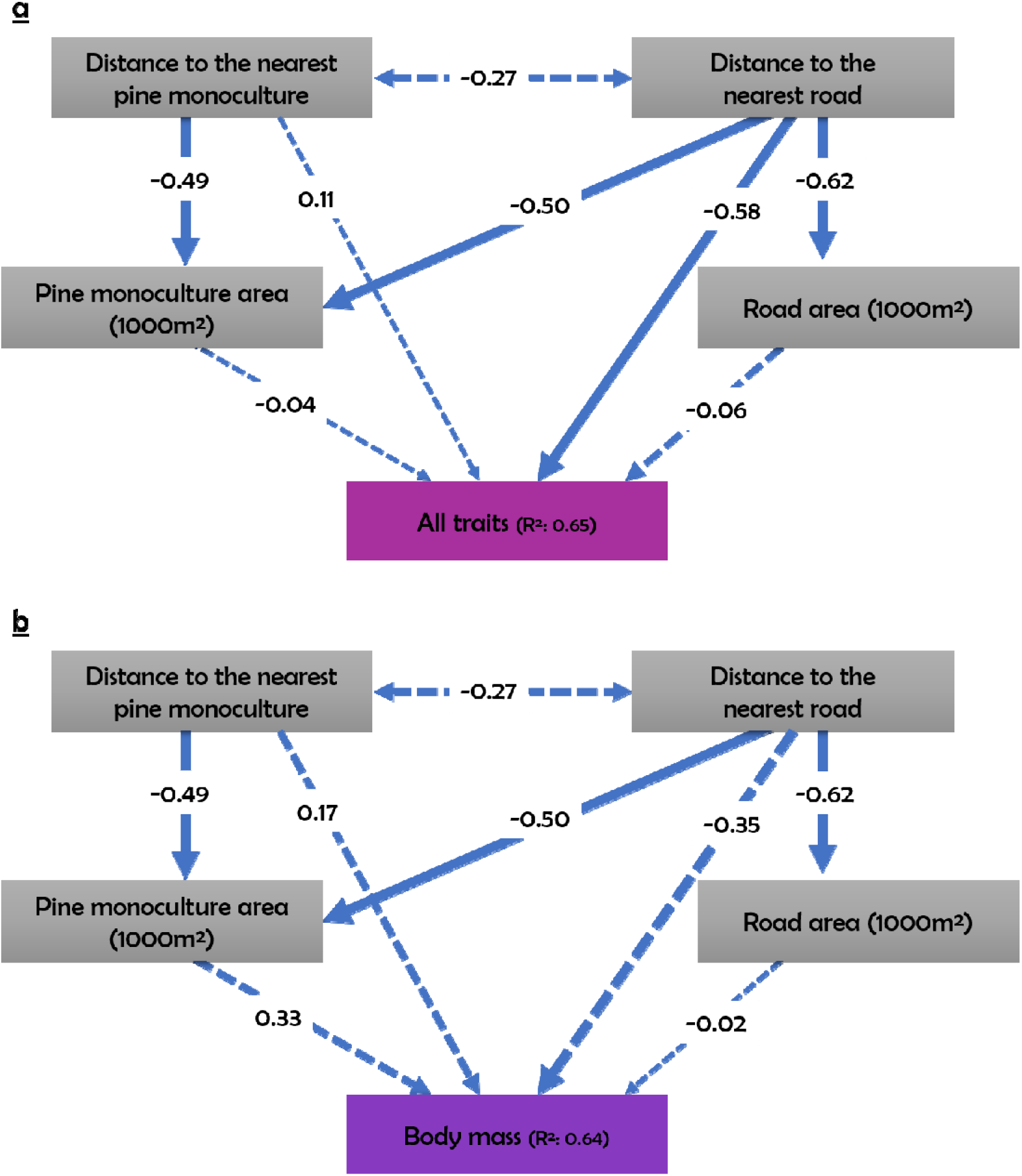
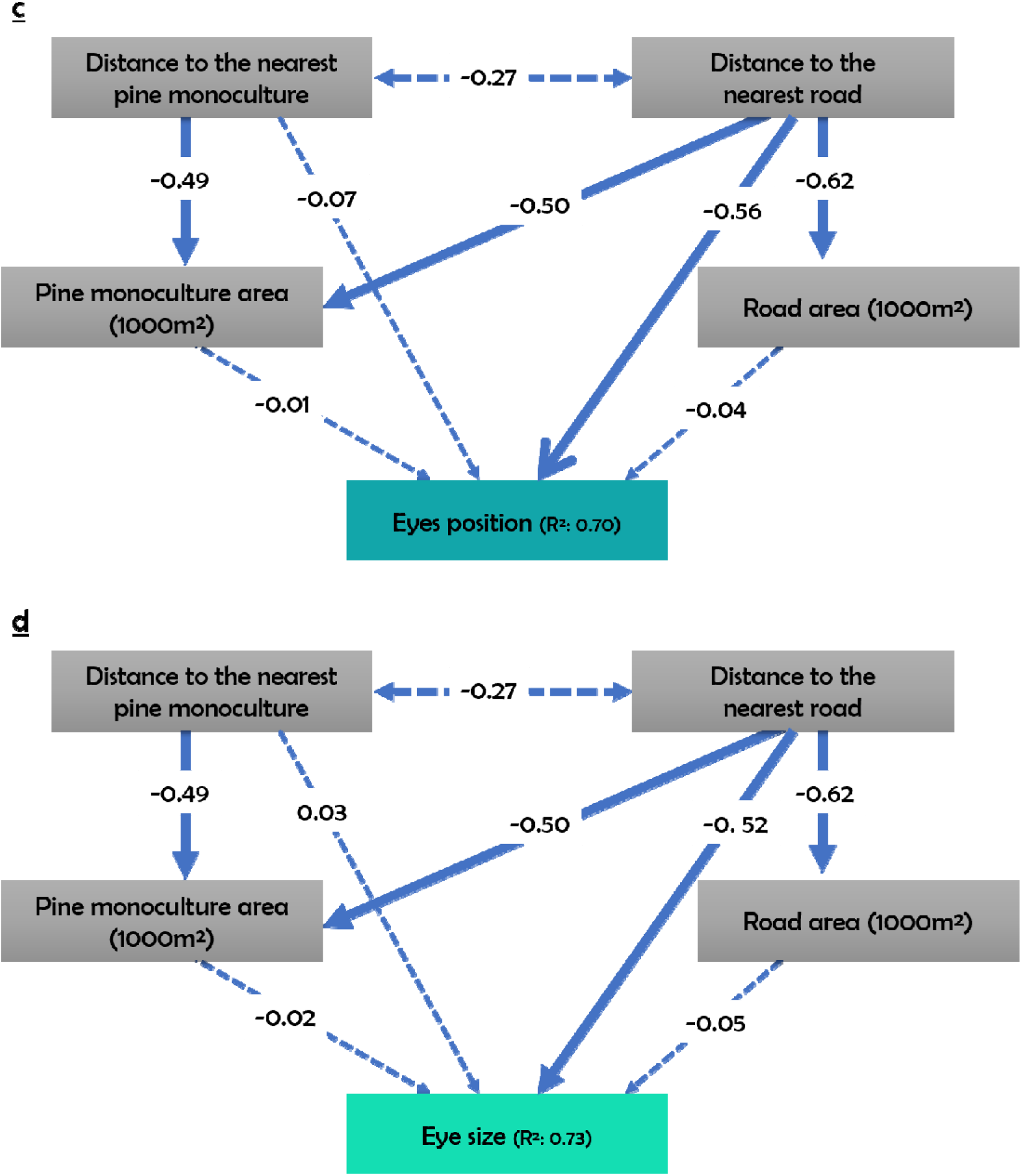
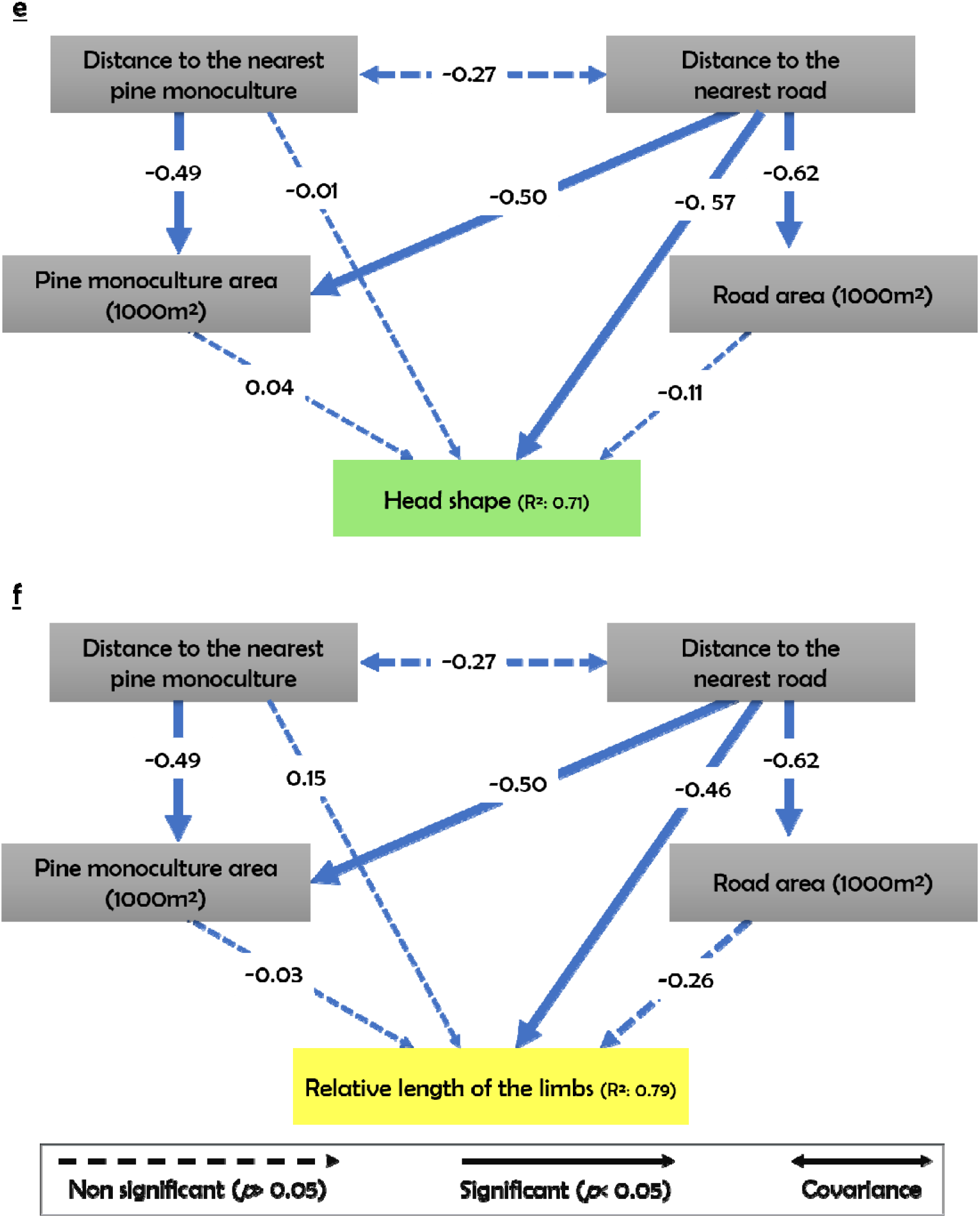
Schematic structural equation models showing the pathways by which the anthropogenic stressors affect the whole of the analysed functional traits – a), all functional traits of the anuran metacommunity; b) body mass; c) eye position; d) eye size; e) head shape; f) relative length of the limbs.

## Discussion

In this study we show that functional and phylogenetic compositions of an anuran metacommunity in a subtropical environment are affected by two anthropogenic stressors – roads and pine monocultures –, although the weight of these relations varies with each compositional facet evaluated. Distance to roads affects mosty the functional composition, while the distance to *Pinus* monocultures affects mainly the phylogenetic composition. These results support our initial hypotheses and allow us to infer that functional and phylogenetic components are more suitable to test the influence of anthropogenic stressors on the structure of metacommunities than the taxonomic component (at least for areas affected by the presence of roads and *Pinus* monocultures). Confirming our initial prediction, anthropogenic stressors seem to affect the sets of functional traits differentially, although the functional dispersion of the traits and the functional composition of the anuran metacommunity are influenced by the same stressors; thus, our results demonstrate that the different metrics of functional diversity are consistent in their responses to anthropogenic stressors.

### Distance to roads influences the patterns of functional structure in anuran metacommunities

Habitat modifications by human-induced factors lead to changes in the patterns of metacommunity structure in anurans (Salice et al., 2011; Shanafelt et al., 2018; Dalmolin et al., 2019; Dalmolin et al., 2020). The relative distance to roads influences the functional dispersion and the functional composition of the anuran communities. In general, anuran abundance reduces largely in response to road proximity (Marsh et al., 2017); also, ponds near roads tend to present lower diversity values (e.g. Cosentino et al., 2014). A possible explanation to the decrease in abundance and diversity is that calling activity tends to reduce in breeding sites near roads. In fact, anuran calls may change in frequency and duration in response to traffic noise (Kaiser & Hammers, 2009; Caorsi et al., 2017). As calls are part of the reproductive repertoire of anurans, changes in the acoustic parameters of the reproductive calls in response to the noise cause by traffic may reduce the encounter rate between males and females and, consequently, the reproductive success of several species (Karraker & Gibbs, 2011).

The negative effect of the proximity of ponds to roads on the functional dispersion may suggest that this stressor is acting as a physical barrier that makes it difficult for adults to move between sites (Fahrig et al., 1995; Carr & Fahrig, 2001; Marsh et al., 2017). So, communities close to each other tend to be functionally redundant (Berriozabal-Islas et al., 2018; Leão-Pires, Luiz & Sawaya, 2018; Dalmolin et al., 2019). This effect can be accentuated for many the species of the Bufonidae, which have shorter limbs and move through small jumps, and for the Leptodactylidae, which have limbs specialized for the aquatic environment, which may confer lesser motility in terrestrial environments. The risk of fatality by collision during dispersal from one habitat to another – which may be a water body or a forest fragment – may be high for reproductive individuals of species that have aquatic larvae (Becker et al., 2010), which is precisely the case for all species present in our study area. Still, the tendency is that increased dispersal movements are expected when habitats are devoid of natural vegetation cover and adequate microclimate conditions (Becker et al., 2010), as tends to occur in areas adjacent to roads and *Pinus* monocultures (Dalmolin et al., 2019). This is also supported by the results obtained in the path analysis, especially for the model for the relative length of the limbs. The physical barriers created by roads may also affect the genetic structure of the populations that constitute the communities (Hels & Buchwald, 2001; Langen, Ogden & Schwarting, 2009; Zimmermann Teixeira et al., 2017; Gonçalves et al., 2018). Indeed, when physical barriers are present, inbreeding tends to increase significantly, while heterozygosity is drastically reduced (Hels & Buchwald, 2001). Consequently, cases of malformation in individuals rise, simultaneously to reductions in population size and, subsequently, in community diversity (Gibbs & Shriver, 2005; Cosentino et al., 2014).

The low functional diversity in the communities closer to roads compromises the use of the functional space by the species and, potentially, also the establishment in ecological niches available. For example, head size usually correlates with the mouth opening, associated with bite force and diet diversity in frogs (Emerson, 1985; Emerson & Brumble, 1993; Lappin et al., 2017); consequently, the more diverse the values of these traits, theoretically the greater the occupation of the functional space by the set of species present. However, in less diverse communities, important and available food resources may be lost or become available to other taxa, such as invasive anurans. Recent reviews confirm that invasive species use the breaches in the functional space as gateways to invade communities where they would not occur naturally or establish easily (Lososová et al., 2015; Finerty et al., 2016). For South American frogs this possibility is concrete, as several communities suffer from the invasion of the American bullfrog (*Lithobates catesbeianus*), including communities in the extreme south of Brazil (Both et al., 2014; Both & Melo, 2015). In terms of conservation, this is worrying, as the resilience of communities and the integrity of the ecosystem functions provided by native anurans may be compromised. Indeed, the American bullfrog, for example, is among the world’s 100 most prominent aquatic invasive species causing negative direct and indirect effect on native aquatic fauna worldwide, particularly on native anurans (Bucciarelli et al., 2014), and Brazil is no exception (Both et al., 2014; Both & Melo, 2015).

### Distance to *Pinus* monocultures as the main anthropogenic stressor affecting the patterns of phylogenetic composition in anuran metacommunities

Distance to *Pinus* monocultures affects anuran phylogenetic composition. This may be related to phylogenetic homogenization in ponds near monocultures, as these exotic plantations show much lower plant diversity when compared to native forests (Martínez & Jauregui et al., 2016), as well as there is much lower number of vegetation types within the ponds (Grass et al., 2015); indeed, several of the sampled ponds located within *Pinus* monocultures did not present any kind of vegetation (D.A.D. personal observation).

Exotic plantations – of the genera *Pinus* and *Eucalyptus* – alter soil characteristics through the release of toxic substances, also contributing to lower quality litter (Machado, Moreira & Maltchik, 2012; Ferreira et al., 2015), making these areas inhospitable for anurans dependent on higher soil/water quality or the presence of vegetation for foraging or reproducing. Also, the presence of these forestry monocultures in areas originally formed by grasslands may create physical barriers that hinder the movement of species that do not have specialized morphological traits to climb the vegetation (adhesive discs, for example; Dalmolin et al., 2020). In addition, these monocultures may represent an insurmountable discontinuity between optimal aquatic and terrestrial habitats, which is particularly serious for anurans that depend directly on both habitats (Vasconcelos & Doro, 2016). On the other hand, anurans with higher reproduction rates and rapidly reaching sexual maturity may be less susceptible to road or exotic plantation effects (Grace, Smith & Noss, 2017) and may thrive in the absence of more sensitive species.

Looking at the phylogenetic composition, it becomes apparent that anuran occurrence and distribution in ponds result from ecological-evolutionary relationships affected by recent human-induced changes in the landscape (Jetz & Pyron, 2018; Campos et al., 2019). The main fact to be highlighted here, however, is that even species with wide geographical distribution, and considered to be undemanding in terms of habitat integrity, were susceptible to the presence of monocultures surrounding the ponds where they occur. Habitat loss and modification restrict the occurrence and distribution of neotropical anurans (including generalist species) at different spatial and geographical scales (Vasconcelos & Doro, 2016), which supports the idea that the loss of evolutionary lineages in modified landscapes does not occur at random. Consequently, in modified areas, poorer communities, and most likely representing nested clusters formed from communities less subject to anthropogenic disturbances, are expected (Almeida-Gomes et al., 2019).

### Recommendations for management

Anurans are one of the most diverse and, simultaneously, one of the most threatened vertebrate taxa in the world (Frost, 2011). The rapid transformation of natural habitats into anthropogenic landscapes is an insurmountable reality, especially in Brazil, where the agricultural land and the road network coverage have almost doubled in the last two decades (Zalles et al., 2019). Additionally, much of the infrastructure expansion is towards the proximity of protected areas, which will certainly compromise the functionality and effectiveness of these areas for conservation (Coelho et al., 2012).

Here we showed that roads and exotic monocultures significantly affect the structuring of anuran metacommunities. We underline the importance of ensuring high soil and water quality within and around ponds, and of maintaining buffers of native vegetation and their connectivity in the surrounding areas, guaranteeing the existence of gradients of environmental heterogeneity in both the vegetation and the substrate (McKinney, 2002; Lion et al., 2014; Berriozabal-Islas et al., 2018; Hansen et al., 2019). These environmental variables proved to influence the structure of tropical anuran communities (da Silva, Candeira & Rossa-Feres, 2012; Prado & Rossa-Feres, 2014; Dalmolin et al., 2019) and have a fundamental role in the intra and interspecific functional variation that may be necessary for species to resist in these systems (Dalmolin et al., 2020). In addition, the construction of road ditches or artificial ponds close to the roads, as well as the use of attractive lamps in energy networks (which end up attracting insects), should be avoided in order not to attract anurans to these areas where the risk of being run over is high (Coelho et al., 2012). Finally, mitigation actions could involve controlling the flow of vehicles at times coinciding with the reproductive season of the majority of the species present (mainly Austral spring and summer; Coelho et al., 2012; Eberhardt, Mitchell & Fahrig, 2013). Such measures are rather straightforward, particularly in protected areas, as is the case of the PNLP, and should be able to increase anuran species’ and assemblages’ resilience, compensating for the energetic demands resulting from the physiological adjustments required to inhabit the suboptimal habitats created by anthropogenic modifications of the landscape (Morley et al., 2019; Rivera-Ordonez et al., 2019; Rubalcaba, Gouveia & Olalla-Tárraga, 2019). These measures should also favour dispersal between ponds, increasing the chances of colonization of other assemblages and, consequently, promoting gene flow (Rittenhouse, Semlitsch & Thompson, 2009).

## Acknowledgments

We thank Conselho Nacional de Desenvolvimento Cienti’fico e Tecnolo’gico (CNPq) for financial and logistical support and for the Diego Dalmolin’ Phd scholarship and Maria João Pereira’ CNPq Research Productivity scholarship. We also thank Lagoa do Peixe National Park for all the support during the field activities. This study was done under ICMBIO licence number 55409–1. Finally, we thank all those who helped during field and lab work and the Leandro Duarte (UFRGS), Luís Bini (UFG), Márcio Borges Martins (UFRGS) and Vinicius Bastazini (CNRS) for their contributions to previous versions of this manuscript.

